# Microsecond Melting and Revitrification of Cryo Samples – Protein Structure and Beam-Induced Motion

**DOI:** 10.1101/2022.02.14.480378

**Authors:** Oliver F. Harder, Jonathan M. Voss, Pavel K. Olshin, Marcel Drabbels, Ulrich J. Lorenz

## Abstract

We have recently introduced a novel approach to time-resolved cryo-electron microscopy (cryo-EM) that involves melting a cryo sample with a laser beam to allow protein dynamics to briefly occur in liquid, before trapping the particles in their transient configurations by rapidly revitrifying the sample. With a time-resolution of just a few microseconds, this approach is notably fast enough to study domain motions that are typically associated with the activity of proteins, but which have previously remained inaccessible. Here, we add crucial details to the characterization of our method. We show that single-particle reconstructions of apoferritin and cowpea chlorotic mottle virus (CCMV) from revitrified samples are indistinguishable from those in conventional samples, demonstrating that melting and revitrification leaves the particles intact and that they do not undergo structural changes within the spatial resolution afforded by our instrument. We also characterize how rapid revitrification affects the properties of the ice, showing that revitrified samples exhibit comparable amounts of beam-induced motion. Our results pave the way for microsecond time-resolved studies of the conformational dynamics of proteins and open up new avenues to study the vitrification process and address beam-induced specimen movement.

**Synopsis:** Microsecond melting and revitrification of cryo samples preserves the structure of embedded particles. The beam-induced motion of revitrified samples is comparable to that of conventional cryo samples.

## 1. Introduction

Cryo-electron microscopy has been undergoing a stunning development in recent years, fueled by the introduction of a number of crucial innovations. Improved sample preparation methods (Russo & Passmore, 2014; Naydenova *et al*., 2020; Dandey *et al*., 2020; Ravelli *et al*., 2020), ever more powerful electron microscopes (Nakane *et al*., 2020; Yip *et al*., 2020), faster and more sensitive electron detectors (Li *et al*., 2013), and more sophisticated computational tools (Punjani *et al*., 2017; Zivanov *et al*., 2020) have recently made it possible to obtain reconstructions of proteins at atomic resolution (Nakane *et al*., 2020; Yip *et al*., 2020). At its current rate of growth, cryo-EM is predicted to rival x-ray crystallography as the most popular method in structural biology in just a few years (Hand, 2020; Callaway, 2020). Another exciting frontier in cryo-EM is the study of dynamics, which can reveal insights into the function of a protein beyond the information that is available from a static structure (Frank, 2004; Chen & Frank, 2016; Frank, 2017). Conformational sorting with advanced computation tools can be used to map out the free energy surface that a protein explores under equilibrium conditions in great detail (Schwander *et al*., 2014; Dashti *et al*., 2014; Zhong *et al*., 2021). However, short-lived intermediates or transient states as well as fast out-of-equilibrium processes are more difficult to access. Time-resolved cryo-EM enables the study of such processes in principle, where dynamics are typically initiated with a light pulse (Menetret *et al*., 1991; Shaikh *et al*., 2009) or through rapid mixing (Lu *et al*., 2009; Shaikh *et al*., 2014; Berriman & Unwin, 1994). The sample is plunge frozen as the dynamics occur, trapping intermediates that can be subsequently imaged (Chen & Frank, 2016; Frank, 2017; Berriman & Unwin, 1994; Unwin & Fujiyoshi, 2012). However, the time-resolution of this approach is several milliseconds and is fundamentally limited by the timescale of plunge freezing, about 1 ms (Frank, 2017; Shaikh *et al*., 2009). It is therefore too slow to capture many relevant processes, in particular the domain motions of proteins that are typically associated with their activity and that frequently occur on the microsecond to millisecond timescale (Henzler-Wildman & Kern, 2007; Boehr *et al*., 2006).

We have recently established a novel approach to time-resolved cryo-EM that affords microsecond time-resolution (Voss *et al*., 2021*a,b*). We illuminate a cryo sample with a focused laser beam to locally melt it and thus allow particle dynamics to briefly occur in liquid. After tens of microseconds, the heating laser is turned off and the sample rapidly cools and revitrifies, arresting the particles in their transient configurations, in which they can be subsequently imaged. Importantly, the time-resolution of our approach is determined by the timescale of vitrification, which occurs within just a few microseconds (Voss *et al*., 2021*b*). Our method thus affords a time-resolution that is three orders of magnitude higher than in conventional time-resolved experiments (Shaikh *et al*., 2009; Frank, 2017), which makes it possible to study a wide range of fast dynamics that have previously remained inaccessible.

Here, we add crucial details to the characterization of our method. While we have previously shown that the melting and revitrification process leaves the proteins intact (Voss *et al*., 2021*a,b*), we here corroborate this finding by demonstrating that reconstructions obtained from conventional and revitrified cryo samples are indistinguishable within the spatial resolution of our instrument. Moreover, we analyze how the revitrification process alters the properties of the ice, showing that revitrified samples exhibit similar amounts of beam-induced motion as conventional samples, albeit with small differences in the drift behavior.

## 2. Methods

**Figure 1a,b** illustrates the sample geometry and experimental approach. Cryo samples are prepared on UltrAuFoil R1.2/1.3 300 mesh gold grids (Russo & Passmore, 2014) (see **Note S1**), and the melting laser (532 nm, 24±1 µm spot size in the sample plane, see **Note S2**) is centered on a grid square (**Figure 1a**). Under illumination with a 20 µs laser pulse, the sample rapidly melts in the vicinity of the laser focus and subsequently revitrifies, arresting the motions of the embedded particles (**Figure 1b**).

**Figure 1.**
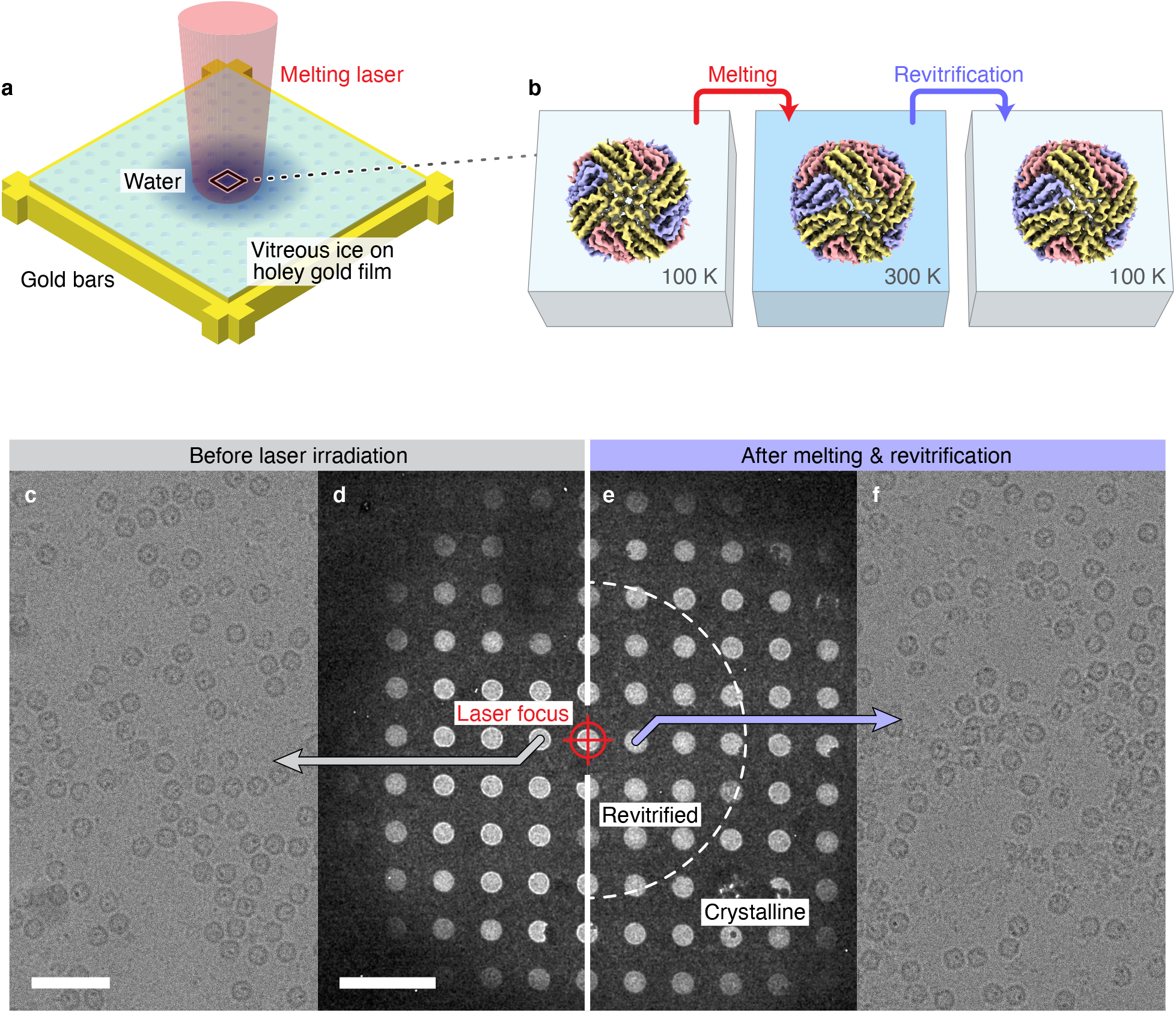
Rapid melting and revitrification of cryo samples, concept and experimental demonstration. (**a**) Illustration of the geometry of the cryo sample, which is prepared on a holey gold film supported by a gold mesh. The sample is irradiated *in situ* with a laser beam that is centered onto a grid square. (**b**) In the vicinity of the laser focus, the sample rapidly melts, allowing embedded particles to undergo equilibrium dynamics in the liquid phase. When the laser is switched off, the sample rapidly revitrifies, trapping the particles, so that they can be subsequently imaged. (**c**–**f**) Micrographs of a cryo sample of apoferritin (**c**,**d**) and of the same sample after melting and revitrification with a 20 µs laser pulse (**e**,**f**). The laser focus is aligned to the central hole, which is marked with a crosshair. The outline of the revitrified area is indicated in (**e**) with a dashed semicircle. Adjacent regions have crystallized. Scale bars, 50 nm in (**c**) and 5 µm in (**d**).

Experiments are carried out with a JEOL 2200FS transmission electron microscope that we have modified for time-resolved experiments (Olshin *et al*., 2020) and that is equipped with a Gatan K3 direct electron detector (**Note S2**). **Figure 1c–f** displays micrographs of a typical experiment with a cryo sample of mouse apoferritin (experimental details in **Note S1**). **Figure 1d** shows a low magnification view of the grid square before laser irradiation, and **Figure 1c** a micrograph collected from the hole marked with a grey arrow. The sample is then illuminated *in situ* with a 20 µs laser pulse (205 mW), with the laser beam aligned to the central hole, which is marked with a crosshair. As indicated in **Figure 1e**, a circular area around the center of the laser focus has melted and revitrified (dashed semicircle). In contrast, the adjacent regions, in which the temperature has remained below the melting point of water, have crystallized (Voss *et al*., 2021*a*). We then collect micrographs from holes within the revitrified area, such as that of **Figure 1f**, which reveals intact apoferritin particles. We note that in order to ensure a comparable temperature evolution in all melting and revitrification experiments, the laser power (typically about 165 mW) was always adjusted to obtain a revitrified region of similar size to that in **Figure 1e** (Voss *et al*., 2021*a*).

## 3. Results and Discussion

Single-particle reconstructions confirm that the melting and revitrification process preserves the structure of apoferritin (Wu *et al*., 2020) (**Notes S3**–**5**). **Figure 2a** compares the density map obtained from a conventional cryo sample (left, 4.60 Å resolution) with that obtained after melting and revitrification (right, 4.25 Å resolution). Individual helices are clearly resolved in both reconstructions. As evident in the details shown on the left and right, some side chain density is visible, which is slightly more pronounced in the higher resolution map obtained from the revitrified samples. Within the resolution afforded by our instrument, the structures are indistinguishable. This result is confirmed by a second set of experiments on cryo samples of CCMV, an icosahedral plant virus (Speir *et al*., 1995; Vriezema *et al*., 2005) (**Figure 2b**). We obtain near-identical reconstructions of CCMV with a resolution of 5.03 Å and 5.20 Å for the conventional (left) and the revitrified cryo samples (right), respectively. Secondary structural elements of the viral capsid are well resolved, which is also evident in the details of the density in the vicinity of the quasi three-fold symmetry axis. We note that the viral RNA inside the capsid is disordered and therefore not resolved. We conclude that the melting and revitrification process leaves the particles intact and that within the spatial resolution of our experiment, it does not alter their structure.

**Figure 2.**
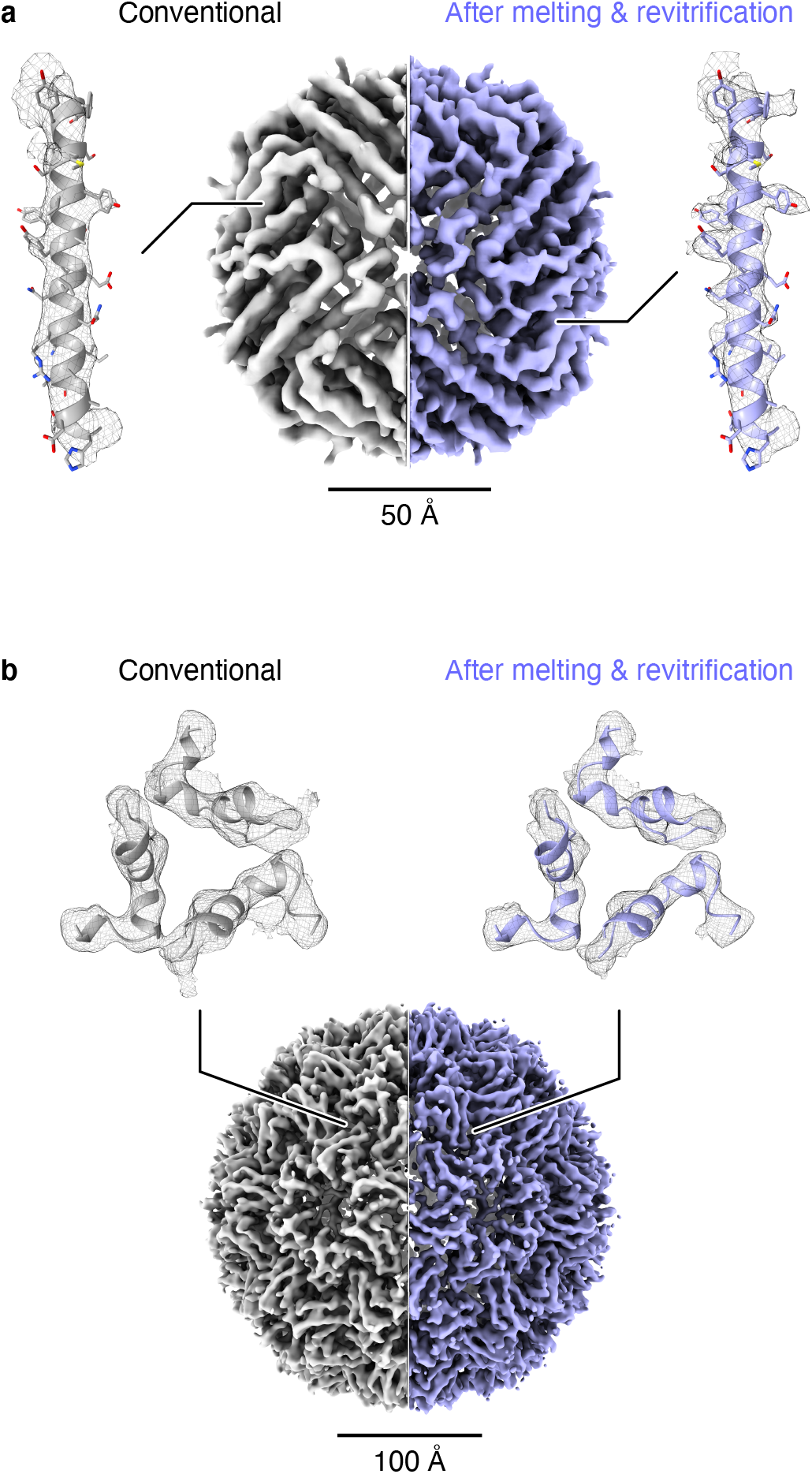
Comparison of single-particle reconstructions obtained from conventional and revitrified cryo samples. (**a**) Reconstructions of apoferritin from a conventional cryo sample (left, 4.60 Å resolution) and a melted and revitrified cryo sample (right, 4.25 Å). Details are shown for the densities of an alpha helix that has been fitted with PDB model 6V21 (Wu *et al*., 2020). (**b**) Single-particle reconstructions of CCMV from a conventional cryo sample (left, 5.03 Å) and a melted and revitrified cryo sample (right, 5.20 Å). Details of the densities of the capsid are shown in the vicinity of the quasi three-fold symmetry axis. The densities have been fitted with PDB model 1CWP (Speir *et al*., 1995). The structures from conventional and revitrified samples are indistinguishable within the resolution of our instrument.

Evidently, rapid melting and revitrification does not expose the particles to any mechanical forces or other processes that damage their structure. This conclusion is supported by a consideration of the different elements of the experiment, including the nature of the interaction between the laser beam and the particles, the melting and revitrification steps, as well as the interactions that the particles encounter while the sample is liquid.

The interaction of the particles with the 532 nm laser beam is only indirect, since neither the particles nor the vitreous ice film absorb visible light, which prevents any type of photodamage. Instead, the laser heats the holey gold film of the specimen grid, which then melts the vitreous ice through fast heat diffusion. We have previously shown that the sample temperature can be controlled to prevent heat denaturation of the particles. In particular, evaporative cooling provides a negative feedback that stabilizes the sample temperature (Voss *et al*., 2021*a,b*).

While the sample is liquid, the particles may interact with the interface between the water film and the vacuum of the microscope. Interactions with the air-water interface have previously been shown to lead to particle denaturation (D’Imprima *et al*., 2019). However, the number of such collisions on the timescale of our experiment, tens of microseconds, is significantly lower than during the plunge-freezing process, where the time between blotting and vitrification is on the order of a second. Furthermore, it has previously been concluded that during this time, a layer of unraveled proteins forms at the aqueous surface that protects particles from reaching the interface (Yoshimura *et al*., 1994; Glaeser, 2018). We therefore expect the number of collisions that lead to denaturation to be small as long as the thickness of the sample remains large enough.

The rapid revitrification of the sample after the laser pulse should certainly not damage the particles, since it is well established that vitrification preserves the structure of a protein. In fact, the significantly higher cooling rate in our experiment (Voss *et al*., 2021*b*), compared with that typically reached during plunge freezing (Frank, 2017; Shaikh *et al*., 2009), should be better suited to trap the room temperature structure of proteins. The melting step resembles the revitrification process in that it occurs on the same timescale of a few microseconds, just with the temperature evolution reversed (Voss *et al*., 2021*b*). One notable difference, however, is that cubic ice forms during the early stages of laser heating, which only melts once the temperature exceeds 273 K (Voss *et al*., 2021*a*). While the formation of cubic ice crystals could conceivably damage the proteins, this does not appear to be the case. In fact, it has recently been shown that the structure of proteins in devitrified cryo samples is unchanged and that devitrification can even be used to reduce sample drift and improve resolution (Wieferig *et al*., 2021).

While we conclude that the structure of the proteins is not altered, an important related question is whether melting and revitrification changes the properties of the ice. In particular, it is important to establish how it affects the beam-induced motion that occurs during imaging and that limits the obtainable spatial resolution. Such motion arises when mechanical stresses in the vitreous ice, which have built up during plunge freezing, are released under electron irradiation (Russo & Passmore, 2014; Naydenova *et al*., 2020; Wright *et al*., 2006; Brilot *et al*., 2012; Zheng *et al*., 2017; Engstrom *et al*., 2021; Thorne, 2020). **Figure 3a** shows that the average cumulative drift of the apoferritin particles, which is displayed as a function of the electron dose, follows a typical behavior for both the conventional (grey) and revitrified cryo samples (purple, see **Note S6**). They initially drift quickly, before slowing to a lower and constant drift rate after a dose of 10– 20 electrons/Å^2^. The revitrified samples exhibit a slightly larger initial drift, but a smaller asymptotic drift rate. Their average cumulative drift therefore drops below that of the conventional samples after 15 electrons/Å^2^. It is 1.5 Å lower at a dose of 60 electrons/Å^2^, for a total drift of 10 Å. Qualitatively similar, although less pronounced differences are observed for the beam-induced motion of the CCMV samples (**Figure 3b**). The cumulative drift of the rivitrified samples drops below that of the conventional samples after 30 electrons/Å^2^ and is 1.3 Å lower at a dose of 60 electrons/Å^2^, for a total drift of 15 Å.

**Figure 3.**
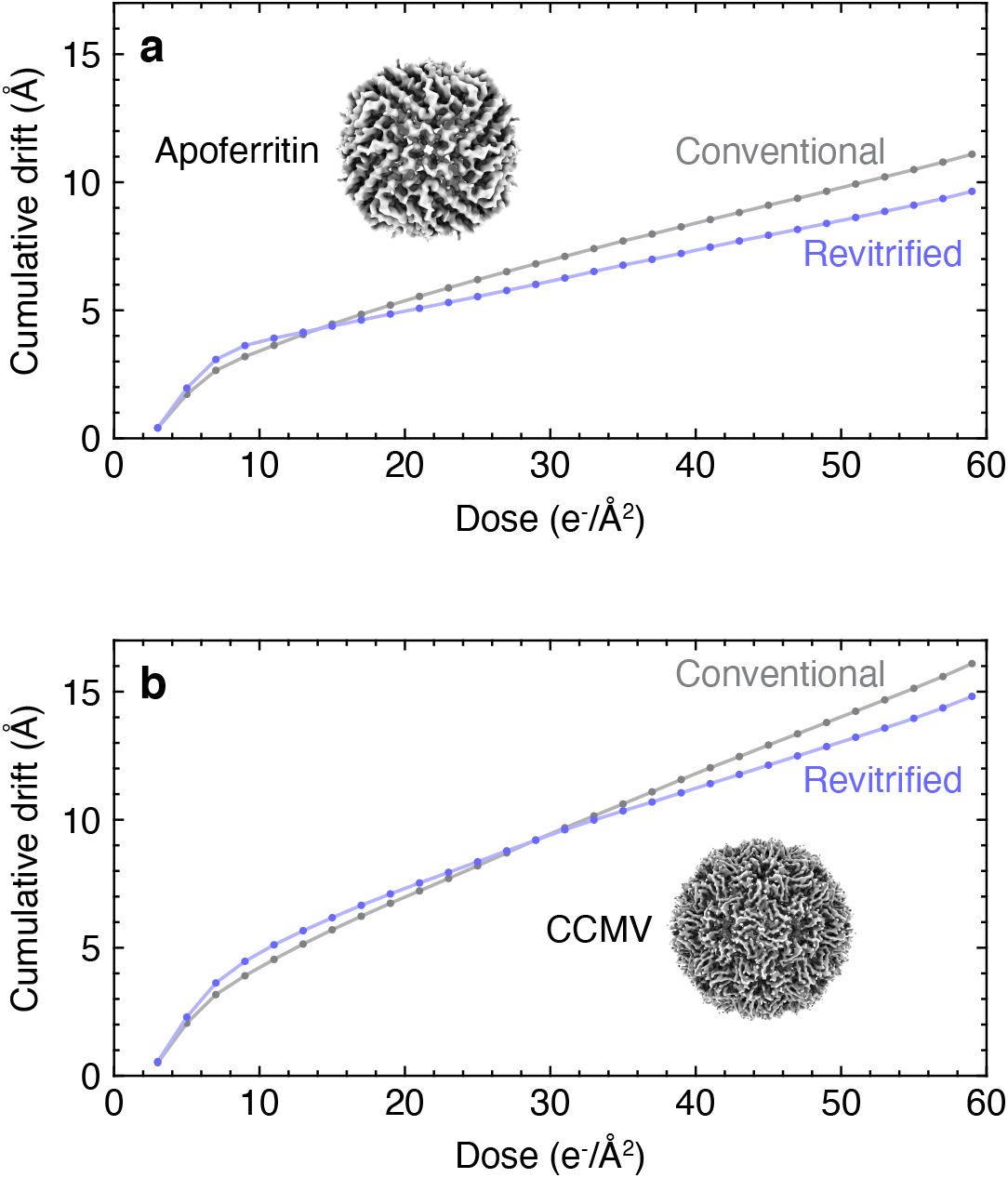
Comparison of sample drift in conventional and revitrified cryo samples. (**a**) Average cumulative specimen drift of apoferritin in a conventional cryo sample (grey) and a melted and revitrified cryo sample (purple). (**b**) Average cumulative specimen drift of CCMV in a conventional cryo sample (grey) and a melted and revitrified cryo sample (purple).

We conclude that revitrified samples exhibit comparable amounts of beam-induced motion to conventional samples, an observation that can shed new light on the mechanism that causes stress buildup in the ice during vitrification. It has been proposed that this stress results from the large temperature difference during plunge freezing between the holey gold film and the grid bars. Because of their large heat capacity, the bars cool more slowly and thus exert a compressive force on the vitreous ice film once they contract (Engstrom *et al*., 2021; Thorne, 2020). Since melting and revitrification is highly localized and the grid bars remain at cryogenic temperatures throughout the entire process, this mechanism alone cannot easily explain the beam-induced motion of the revitrified samples (Voss *et al*., 2021*a,b*). Our results are also seemingly at odds with the previous observation that drift increases with vitrification speed, which has been ascribed to the fact that faster cooling provides less time for stresses in the ice to dissipate during plunge freezing (Wu *et al*., 2021; Engstrom *et al*., 2021). Since the cooling rate during revitrification is almost two orders of magnitude higher, one would therefore expect the beam-induced motion to become even more pronounced. We speculate that this is not the case since revitrification removes stresses that arise from large-scale deformations of the entire grid during plunge freezing (Engstrom *et al*., 2021; Thorne, 2020), which compensates for the additional local stress induced by the high cooling rate.

## 4. Conclusion

Our experiments demonstrate that rapid melting and revitrification of cryo samples leaves embedded particles intact, providing a central piece of evidence that our approach is suitable to study the dynamics of proteins on the microsecond timescale. We find that proteins do not undergo structural changes within the spatial resolution afforded by our instrument. However, subtle changes may become apparent at higher resolution, a possibility that we have already begun to explore. Temperature-resolved cryo-EM experiments have established that different conformational ensembles are obtained when the sample is equilibrated at different temperatures before vitrification (Fischer *et al*., 2010; Chen *et al*., 2019). Such structural changes may even occur as the sample is cooled during the plunge-freezing process, depopulating some conformational states that are otherwise present at physiological temperatures (Mehra *et al*., 2020). Melting and revitrification may provide a tool to investigate such effects systematically and even repopulate high-temperature states that are otherwise inaccessible. We also expect that the significantly faster vitrification speed in our experiments should lead to a higher glass transition temperature (Angell, 2008) and thus cause subtle changes in the structure of the water network surrounding the proteins, which should become evident at atomic resolution (Nakane *et al*., 2020; Yip *et al*., 2020).

Our experiments also open up new possibilities for studying the vitrification process in real time (Olshin *et al*., 2021) and how it leads to the buildup of stress that results in beam-induced motion. By suitably modulating the laser power, the cooling rate can be varied by more than three orders of magnitude, and the temperature evolution of the sample can be precisely controlled. This has been difficult to achieve with plunge freezing, where the complex fluid dynamics of the cryogen and its vapor phase as well as the heterogeneous heat transfer properties of the sample lead to widely varying vitrification speeds, even across a single specimen grid (Wu *et al*., 2021). Even though we observe only small differences in the beam-induced motion of revitrified samples, our initial results suggest that it may be possible to release stress through irradiation with a sequence of laser pulses.

## Supporting Information

Notes for sample preparation, instrumentation, imaging for single-particle cryo-EM, imaging processing and single-particle reconstructions, visualization of reconstructions, and analysis of beam-induced motion.

## Data Availability Statement

The data that support the findings of this study are available from the corresponding author upon reasonable request.

## Acknowledgements

We kindly thank Prof. Henning Stahlberg and Dr. Dongchun Ni (LBEM EPFL, Switzerland) as well as Dr. Kelvin Lau (PTPSP EPFL, Switzerland) for providing the apoferritin samples. We gratefully acknowledge Prof. Jeroen Cornelissen and Dr. Regine van der Hee (BNT University of Twente, The Netherlands) for providing the CCMV samples. We would also like to thank the Interdisciplinary Centre for Electron Microscopy at EPFL for access to sample preparation equipment, and we kindly acknowledge Dr. Sergey Nazarov (DCI Lausanne, Switzerland) for helpful discussions regarding single-particle reconstructions. This work was supported by the ERC Starting Grant 759145 and by the Swiss National Science Foundation Grant PP00P2_163681.

