## Supporting Information for "Microsecond Melting and Revitrification of Cryo Samples – Protein Structure and Beam-Induced Motion"

**Note S1. Sample preparation**

Cryo samples were prepared on UltrAuFoil R1.2/1.3 300 mesh grids (Quantifoil). The grids were rendered hydrophilic through 10 minutes of plasma cleaning in an ELMO glow discharge system operating with negative polarity, 0.8 mA plasma current, and 0.2 mbar residual air pressure. Approximately 3–3.5 µl of sample solution were applied to the foil side of the grids – either 1.70 mg/ml mouse heavy chain apoferritin in 20 mM HEPES buffer with 300 mM sodium chloride at pH 7.5 (Figure 1), 0.33 mg/ml mouse heavy chain apoferritin in 20 mM Tris buffer with 150 mM sodium chloride at pH 7.4 (Figure 2,3), or 11.5 mg/ml CCMV in 0.1 M sodium acetate buffer with 1 mM Na2EDTA and 1 mM NaN3 at pH 5.0 (Figure 2,3). The samples were plunge frozen with a Vitrobot MarkIV (Thermo Fisher Scientific) that was operated at 100 % relative humidity and a temperature of 22 °C, using a blotting time of 3–4 s. For imaging, the grids were loaded into a single tilt cryo-transfer specimen holder.

**Note S2. Instrumentation**

Experiments were performed with a JEOL 2200FS transmission electron microscope that we have modified for time-resolved experiments, as previously described (Olshin *et al.*, 2020). The microscope is operated at 200 kV accelerating voltage and is equipped with an in-column Omega-type energy filter and a K3 direct electron detector (Gatan). Microsecond laser pulses for *in situ* melting and revitrification of the sample are obtained by chopping the output of a continuous laser (532 nm) with an acousto-optic modulator. The laser is reflected off a mirror located above the upper pole piece of the objective lens and strikes the sample at close to normal incidence. The laser beam is focused to a spot size of 24±1 µm FWHM, as measured by a knife-edge scan in the sample plane.

**Note S3. Imaging for single-particle cryo-EM**

Micrographs were zero-loss filtered with a 10 eV slit width. The electron detector was operated in counting mode to acquire 30-frame movies with a total dose of 60 electrons/Å$^2$ and 2 and 3 s exposure times for apoferritin and CCMV, respectively. Micrographs were recorded with a pixel size of 0.9750 Å and 0.8514 Å for apoferritin and CCMV, respectively. Defocus values were in the range of 0.5–2.5 µm.

**Note S4. Image processing and single-particle reconstructions**

All images were corrected for the magnification distortion (Grant & Grigorieff, 2015), which we determined from micrographs of a gold calibration standard (Ted Pella, product #673) with the mag_distortion_estimate_1.0.1 script (Grant & Grigorieff, 2015). Single-particle reconstructions were carried out with cryoSPARC version 3.2.0 (Punjani *et al.*, 2017). The conventional apoferritin dataset consisted of 167 images. After patch motion correction and patch CTF estimation, 109 micrographs with CTF fits below 6.5 Å were selected. Following blob picking and inspection, particles were sorted into 30 classes, and 5 were used as templates to pick 146,763 particles from the micrographs. Of the picked particles, 81,332 were selected and sorted into 50 classes with a 160 Å circular mask. After removal of junk, 76,144 particles from 38 classes were selected and subjected to another round of 2D classification (50 classes, 140 Å mask). From this round of sorting, 55,183 particles across a total of 13 classes were selected. After *ab initio* reconstruction (O symmetry) and heterogeneous refinement (O symmetry) into 2 classes, the high-resolution class (49,885 particles) was homogeneously refined using O symmetry, resulting in a 4.60 Å map.

The revitrified apoferritin dataset was comprised of 100 micrographs. After patch motion correction and patch CTF estimation, 70 images with CTF fits under 5.5 Å were chosen. Following blob picking and inspection, particles were sorted into 30 classes, and 27 used as templates to pick 93,800 particles from the micrographs. Next, 58,792 particles were selected and sorted into 50 classes with a 160 Å circular mask. After removal of junk, 53,783 particles from 33 classes were chosen and again subjected to another round of 2D classification (50 classes, 140 Å mask). A total of 40 classes, corresponding to 49,476 particles, were selected. After *ab initio* reconstruction (O symmetry) and heterogeneous refinement (O symmetry) into 2 classes, the high-resolution class (45,953 particles) was homogenously refined using O symmetry, resulting in a 4.25 Å map.

The conventional CCMV dataset consisted of 269 images. After patch motion correction and patch CTF estimation, 97 micrographs with CTF fits below 6.0 Å were selected. Following blob picking and inspection, particles were sorted into 30 classes, and 27 were used as templates to pick 36,783 particles from the micrographs. Of the picked particles, 6,616 were selected and sorted into 50 classes with a 320 Å circular mask. After removal of junk, 6,554 particles from 21 classes were selected. After *ab initio* reconstruction (**I** symmetry) and heterogeneous refinement (**I** symmetry) into 2 classes, the high-resolution class (3,560 particles) was homogeneously refined using **I** symmetry, resulting in a 5.03 Å resolution map.

The revitrified CCMV dataset consisted of 391 images. After patch motion correction and patch CTF estimation, 120 images with CTF fits below 6.5 Å were chosen. After blob picking and inspection, particles were sorted into 30 classes, and 10 were used as templates to pick 42,201 particles from the micrographs. Of the picked particles, 7,866 particles were selected and sorted into 50 classes with a 320 Å circular mask. After removal of junk, 7,240 particles from 16 classes were selected. After *ab initio* reconstruction (C1 symmetry) and heterogeneous refinement (**I** symmetry) into 2 classes, the high-resolution class (5,029 particles) was homogeneously refined using **I** symmetry, resulting in a 5.20 Å resolution map.

**Note S5. Visualization of reconstructions.**

Single-particle reconstructions were visualized with ChimeraX (Goddard *et al.*, 2018). The volumes of apoferritin in **Figure 2** are displayed with contour levels of 0.20~~0~~ and 0.15~~0~~ for the conventional and revitrified samples, respectively. The molecular model from PDB 6V21 (Wu *et al.*, 2020) was placed into the density through rigid body fitting. The details in **Figure 2** show the density within 3.3 Å of residues 13–42 of chain C. The volumes for CCMV are displayed with contour levels of 0.100 and 0.115, for the conventional and revitrified samples, respectively. The molecular model from PDB 1CWP (Speir *et al.*, 1995) was placed into the density through rigid body fitting. Details are shown for the density within 3.3 Å of residues 76–86 and 142–153 of chains A, B, and C.

**Note S6. Analysis of beam-induced motion**

Drift trajectories of apoferritin and CCMV were determined with local motion correction in cryoSPARC. The average displacement of the particles as a function of dose (**Figure 3**) was calculated using only the particles that were included in the reconstructions in **Figure 2**. We note that if we include all particles, the result is qualitatively the same.
